# Bioenergetic metabolism restarts alongside germinant sensing and hydration in bacterial spore germination

**DOI:** 10.1101/2025.03.13.642030

**Authors:** P Gupta, R Caldbeck, RC Walters, EC Wells, BL Hardman, G Christie, RJ Springett, JN Blaza

## Abstract

Bacterial spore germination transforms spores from dormant bodies to vegetative cells. Initiation of germination or spore rehydration can proceed without exogenous energy sources, so bioenergetic processes in germination have been overlooked, despite many germinants being energy sources, such as sugars. Here, we apply remission spectroscopy to noninvasively measure the energy-transducing electron-transport chain of intact spores during germination. In *Bacillus megaterium* and *Bacillus subtilis*, we find energisation of cytoplasmic metabolism and the electron transport chain occurs early in germination, before or alongside rehydration. The *aa*_3_-type oxidases (Qox, Cta) accumulate nonradical ferryl intermediates of their catalytic cycle demonstrating that the electron-transport chain is operating in a regime of high membrane potential. The Yth isoform of the *bd* oxidase is found in abundance in both organisms, functioning to allow rapid electron transfer to O_2_ when the *aa*_3_-type oxidases are hindered, establishing a new role for this enzyme beyond O_2_-scavenging. Deletion of Yth slows rehydration, directly linking bioenergetic processes to germination. Our data lead us to propose a powered germination model, where the Ger-mediated signalling cascade and bioenergetic processes occur simultaneously and semi-independently.

## Introduction

Bacteria of the phylum Firmicutes form endospores as part of their lifecycle. Resistant to desiccation, irradiation, heat, and extremes of pH, these structures are amongst the most robust biological forms. Bacteria sporulate as a survival strategy: once formed, they can maintain viability for prolonged periods of time, allowing them to wait out unfavourable conditions (Kennedy et al., 1994). When conditions conducive for growth are re-established, spores sense energy-carrying molecules such as sugars, amino acids, nucleosides, or non-nutrient signals (e.g. K^+^) as ‘germinants’ to initiate germination (Ross and Abel-Santos, 2010). Remarkably, during germination they can transition from being an inert object to a metabolically-active cell in just over an hour (Keijser et al., 2007).

Mature *Bacillus* spores have an onion-like multi-layered architecture that underpins their exceptional robustness (Setlow and Christie, 2023). The DNA-containing core contains precipitated calcium-dipicolinic acid chelate (CaDPA) at >800 mM (Kong et al., 2012). The core has a gel-like consistency with little mobile water (Kaieda et al., 2013; Sunde et al., 2009). Around the core is a cytoplasmic membrane, which harbours the germinant receptors (GRs) and SpoVA channel proteins that are directly involved in the initiation of germination (Paidhungat and Setlow, 2001; Vepachedu and Setlow, 2005), along with respiratory enzymes of the electron-transport chain (ETC) (Chen et al., 2019; Racine and Vary, 1980; Wilkinson and Ellar, 1975). The cytoplasmic membrane presents a permeability barrier, where the constituent lipids are mostly immobilised (Cowan et al., 2004). Around the cytoplasmic membrane is the germ cell wall, peptidoglycan-based cortex, outer membrane, proteinaceous coat, and an exosporium layer in some species. Germination has been described in two stages (Christie and Setlow, 2020; Moir and Cooper, 2015). In stage I, germinants bind to their receptors and there is substantial efflux of H^+^, K^+^, and Na^+^ out of the spore in addition to slow leakage of CaDPA (Wang et al., 2015). As more CaDPA from the spore core is released and replaced by water (rehydration), the spore moves into stage II, where cortex-lytic enzymes are activated allowing spore swelling, emergence, and elongation followed by the first cell division (Ishikawa et al., 1998; Riyami et al., 2019). These later processes are termed outgrowth. Core rehydration occurs during both stage I and II, visualised as phase transition which takes between 2-3 minutes for an individual spore (Pandey et al., 2013; Wang et al., 2015).

Dormant spores are considered energetically depleted. From a bioenergetic viewpoint, cells maintain certain reactions away from equilibrium to create energy-carrying chemical potentials. These reactions are coupled to energetically unfavourable reactions to drive them; the most well-known of these is the phosphorylation of ADP to generate ATP, catalysed by ATP synthase, driven by the imbalance of charge (ΔΨ) and pH (ΔpH) across an energy-transducing membrane. This imbalance of charge is created because the enzymes of the respiratory electron transport chain (ETC) pump protons as they pass electrons from cytoplasmic low-potential donors such as NADH and succinate to high potential acceptors such as O_2_. As the adenine nucleotide pool within dormant spores is largely found as ADP and AMP and the NAD(P)H pool is largely oxidised, these pools are incapable of powering reactions (Ghosh et al., 2015; Setlow and Setlow, 1977; Setlow and Kornberg, 1970). The fact that these pools are not replenished before rehydration begins and that hydration can occur in anaerobic conditions has led to the idea that energy metabolism restarts later in the germination process, even though many germinants are excellent nutrients.

*Bacillus* spores sense the availability of nutrients using germinant receptors. Once the germinants bind to their cognate receptors, the germination cascade is initiated to exit dormancy. Recently, the GerA germinant receptor, ectopically expressed in vegetative cells, was shown to be a membrane channel that conducts cations (Gao et al., 2023). A SpoVAF-FigP channel also conducts cations to amplify the response (Gao et al., 2024). Cation release functions as a biochemical signal but if cation efflux is not matched with anion efflux to cancel charge movement, it will also generate a membrane potential (ΔΨ). Like ATP generation (the phosphorylation potential), ΔΨ is a fundamental mechanism of energy conservation that is linked to many other biochemical reactions, especially active transport and ATP synthesis, making it an attractive hypothetical energy source for the spore. Measurements on single spores have even supported the idea that ΔΨ exists before germination, although this is most likely an artefact stemming from the use of thioflavin-T (ThT), which binds nonspecifically to the spore coat (Kikuchi et al., 2022; Li et al., 2023).

Effective tools for studying molecular events inside spores during germination are lacking. As spores hydrate during germination, the density of their core becomes closer to that of the surrounding medium, so they lose refractivity and can be monitored using transmission absorbance (attenuance) spectroscopy and phase-contrast light microscopy; these approaches have been the bedrock of the field for many decades (Powell and Hunter, 1955). More invasive techniques have heavy caveats to their interpretation. GFP-tagging proteins of interest (Sinai et al., 2015) to track their translation early in germination gives artefactual results when compared to Western blotting and mass-spectrometric measures of protein abundance; the increase in fluorescence intensity is instead attributable to the dramatic physicochemical changes that occur during core rehydration (Swarge et al., 2020b). Exogenous probes are unsuitable because spores have evolved to resist the intrusion of most compounds into their core. In contrast, vegetative cells are typically more susceptible, therefore, along the germination cascade is an axis of increasing permeability. This axis makes it hard to distinguish between real changes occurring during germination and changes to spore structure; the binding of ThT above is one example as are carbocyanin dyes (Magge et al., 2009).

We focus here on two of the most used model spore formers: *Bacillus megaterium* and *Bacillus subtilis*. *B. megaterium* spores rapidly germinate when they sense their potent germinant: glucose. *B. megaterium* was a foundational organism for the germination field (Dills and Vary, 1978; Powell, 1951; Sano et al., 1988; Vary, 1994). *B. subtilis* spores germinate less rapidly, but the extensive molecular tools available and adoption of this organism for sporulation studies has led to much recent work moving to *B. subtilis*.

Here, we use a bioenergetic chamber to make visible-wavelength spectroscopic measurements on intact spores germinating with their favoured germinants: *B. megaterium* on glucose; *B. subtilis* on alanine and glucose. We measure glucose oxidation occurring alongside hydration and ETC reduction before measurable hydration in *B. megaterium*; in *B. subtilis*, rehydration and glucose-powered ETC reduction start simultaneously. These observations challenge models of germination in which metabolism restarts only once core rehydration is complete. We find that the *aa*_3_-type terminal oxidases (CtaA-D & QoxA-D) have different abundances of catalytic intermediates compared to turnover in purified enzymes or vegetative cells, with the nonradical ferryl (F) intermediate predominating, most likely as they are operating in conditions of high ΔΨ created by the efflux of monovalent cations. The appearance of this intermediate from an early time point indicates that ΔΨ builds very quickly. We find that the spores contain a non-proton pumping CN^-^-insensitive *bd*-type oxidase, which while canonically considered an O_2_-scavenging enzyme, is nevertheless abundant in spores cultivated in O_2_-replete conditions. During germination, this oxidase passes electrons to O_2_ as the high ΔΨ slows catalysis by the proton-pumping respiratory complexes leading to reductive pressure. The deletion of this oxidase in both species leads to delayed rehydration kinetics compared to WT, directly linking resumption of bioenergetic processes to the initiation of germination. Based on these findings, we propose a powered germination model in which germinating spores are in fact not energetically depleted and could support active processes much earlier than is currently appreciated.

## Results

### A bioenergetic chamber measures electron transfer to cytochromes and oxygen during spore germination

Haem groups present in the respiratory cytochromes have characteristic visible-wavelength absorbance bands that change as they undergo reduction. These changes were used as spectroscopic handles in our ‘bioenergetic chamber’, which integrates several components around a culture vessel to measure the bioenergetic status of living cells (Fig 1A). Remission spectroscopy uses light back-scattered (’re-emitted’) off the turbid culture instead of transmitted light (Hollis et al., 2003), enabling experiments with dense spore suspensions. The chamber also incorporates an O_2_ optode to simultaneously measure O_2_ consumption rate (OCR) and a gas delivery system that maintains the spore suspension at a constant O_2_ concentration (Kim et al., 2012).

**Figure 1.**
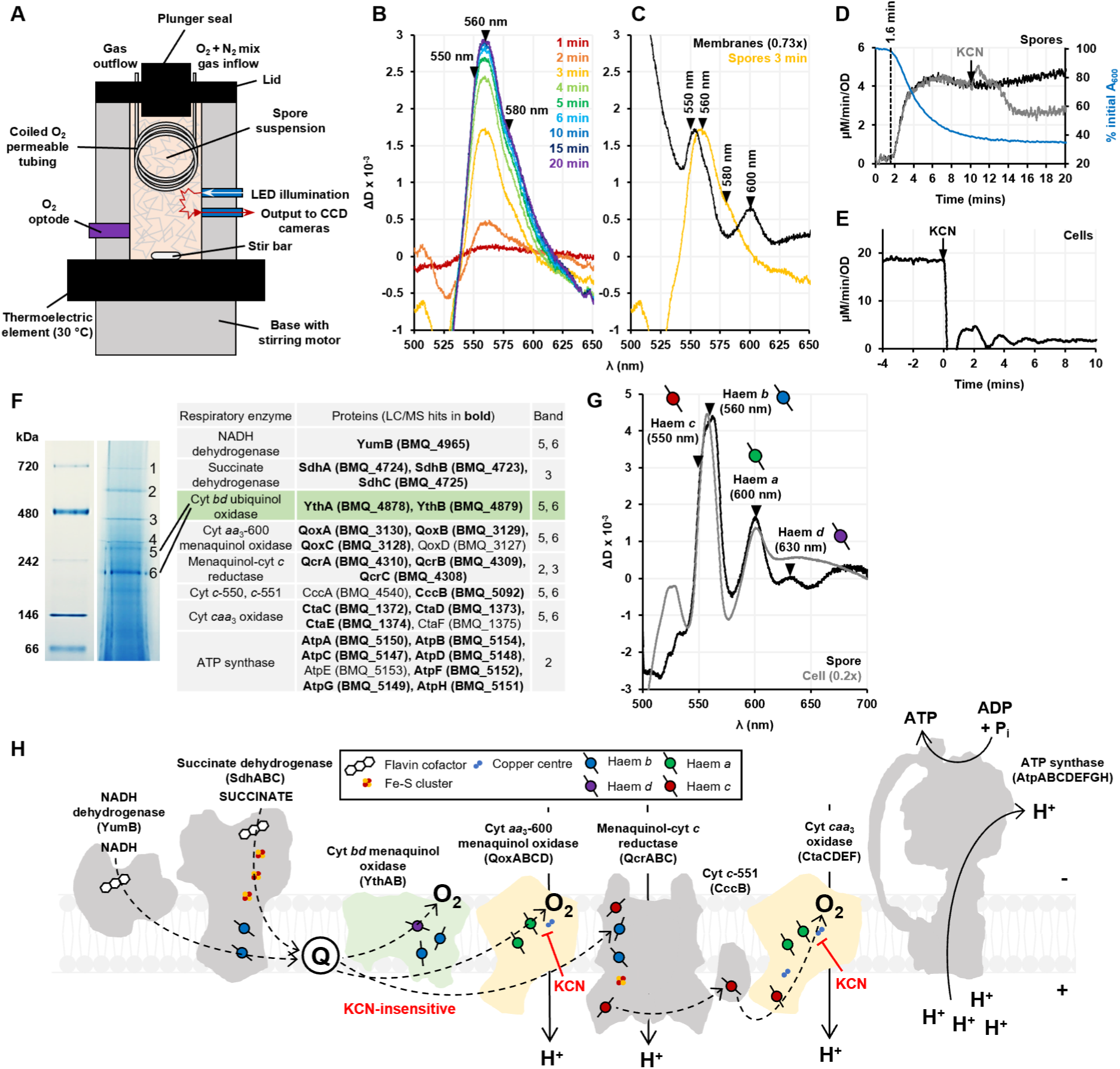
Remission haem spectroscopy on germinating *B. megaterium* spores. (A) The bioenergetic chamber where a dense suspension of heat-shocked spores is maintained at 30 ℃, 120 µM O_2_ and germination is initiated at t=0 min with the addition of 10 mM glucose; spectra are recorded by the CCD-spectrograph system. (B) Difference spectra generated for a given time point (T_p_) after glucose addition at t=0 min. (C) The 3 min spectrum of germinating spores from (B) compared with a scaled spectrum of isolated spore membranes after 3 mins of NADH reduction. Absorbance peaks at indicated wavelengths 550 nm, 560 nm, and 600 nm correspond to reduced haem groups *c*, *b* and *a* respectively, while the broad shoulder at 580 nm is associated with the *aa*_3_-type oxidase F state. (D) OCR of spores germinating with 10 mM glucose (black), poisoned with 1 mM KCN (grey), along with the loss of absorbance rehydration curve (blue). Loss of absorbance starts at 1.6 mins after glucose addition. (E) OCR of late-exponential *B. megaterium* cells grown in LB broth medium inhibited by 1 mM KCN. (F) BN-PAGE gel and the high molecular weight bands labelled 1-6 subjected to LC/MS analysis. The accompanying table lists the proteins of respiratory complexes (along with gene IDs) which were found to be the most abundant in these bands. The cytochrome *bd* paralogue in spores, YthAB, is highlighted in green. (G) The linear regression-subtracted sodium dithionite-reduced minus air-oxidised spectra of isolated cell and spore membranes. (H) Scheme of the ETC present in dormant spores based on the mass spectrometric analysis of BN-PAGE gel bands obtained from DDM-solubilised isolated spore membranes. The CN^-^-sensitive *aa*_3_-type oxidases (Qox and Cta) are shown in yellow and the CN^-^-insensitive *bd* oxidase (Yth) in green. Haem groups and other redox centres are denoted by symbols shown in the legend.

Glucose was added to the suspension of dormant *B. megaterium* spores in the bioenergetic chamber, and spectral changes observed arising from redox changes in the abundant ETC cytochromes (Fig 1B). Overlapping peaks were seen at 550 nm and 560 nm indicating reduction of haem *c* and *b* groups, respectively. Intriguingly, despite the presence of two *aa*_3_-type oxidases (Qox and Cta) in the ETC (Fig 1H), the characteristic signal of haems *a* (peak at 600 nm) was not observed, instead a shoulder was seen at 580 nm. The 580 nm signal is associated with the ferryl intermediate ‘F’ of the *a*_3_-haem in the catalytic cycle of *aa*_3_-type oxidases (Fig 2I). This is a minor intermediate under turnover conditions in purified enzyme preparations (Mason et al., 2014; Rocha and Springett, 2019; Wikström et al., 2023), has been observed in isolated mitochondria and submitochondrial particles (Björck and Brzezinski, 2018; Covian et al., 2023; Shimada et al., 2023; Wikström, 1981), but to the best of our knowledge, this is the first report of the F intermediate being sufficiently abundant to be detectable *in vivo*. Under physiological conditions in mammalian cells, the ferryl intermediates are not detectable, and reduced haem *a* is the dominant spectral signature of cyt *c* oxidase activity (Kim et al., 2011). To understand if this was due to a difference between Qox and Cta in spores and in vegetative cells, we isolated spore membranes, reduced them with NADH and found a characteristic peak at 600 nm, indicative of haems *a* reduction (Fig 1C). We therefore conclude that spore Cta and Qox are canonical *aa*_3_-type oxidases but something within germinating spores places them in a different catalytic state compared to vegetative cells, discussed below.

**Figure 2.**
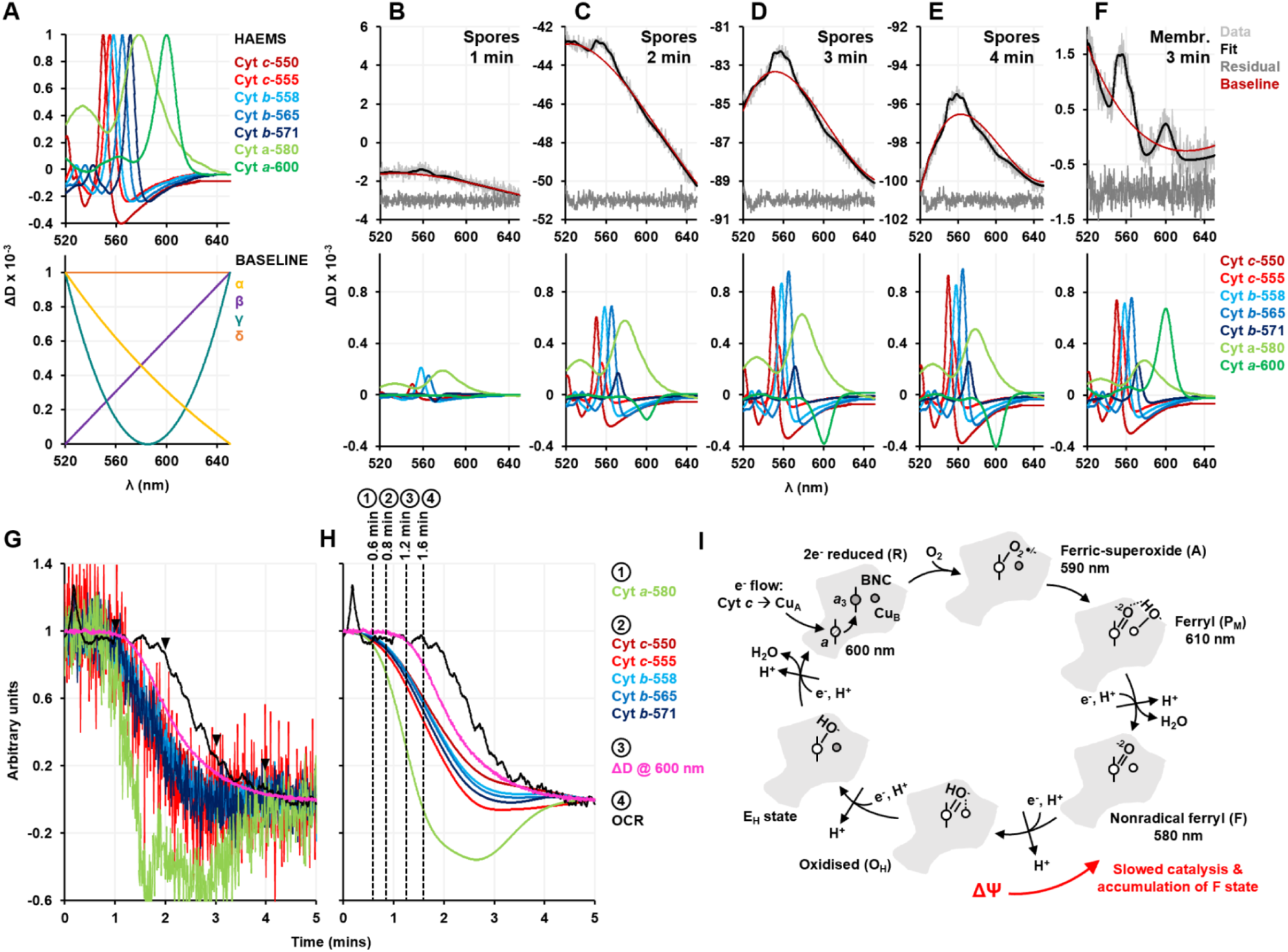
Analysis through spectral unmixing and the kinetics of spectral changes in germinating spores. (A) The haem (top) and baseline (bottom) components of the fitting template. (B)-(E) are spectra of spores germinated with 10 mM glucose, (F) is a spectrum of isolated spore membranes that were reduced with 1 mM NADH. (B)-(F) top panel shows the raw data, the fit imposed, the residuals (suitably offset from 0 to be comparable), and the sum of the baseline components being fitted at the indicated time points. The bottom panel shows the fitting of the haem components. (G) Normalised kinetic traces (fitting versus time) of selected haem components, change in attenuance (ΔD) at 600 nm, and OCR for spores germinated with 10 mM glucose. Arrowheads indicate the time points 1, 2, 3 and 4 mins for which the decomposition analysis is demonstrated in (B)-(E). (H) shows the haem traces smoothed using the Savitzky-Golay filter, along with the unsmoothed rehydration signal and OCR traces. Events in clusters 1, 2, 3, and 4 start 0.6, 0.8, 1.2, and 1.6 mins respectively after glucose addition. (I) The canonical catalytic cycle of *aa*_3_-type oxidases showing the catalytic intermediates formed at the binuclear centre (BNC composed of Cu_B_ and haem a_3_) and the characteristic absorbances of their peaks. Grey colour of the haem/copper group indicates reduced Fe^2+^/Cu^+^, white colour indicates oxidised Fe^3+^/Cu^2+^ state. Haem *a* can be oxidised/reduced, shown as a shaded haem centre in the R intermediate. For every electron and proton transferred to the BNC, another proton is pumped as indicated by the arrows pointing outwards. The text and arrow in red show the effect of ΔΨ on the catalytic cycle.

### The spore electron transport chain is proteomically and functionally distinct from that of a vegetative cell

Given the consensus that bioenergetic processes start later in germination, we were surprised to measure germinating spores consuming O_2_ 1.6 mins after glucose addition, concomitantly with the loss of absorbance at 600 nm (Fig 1D). To understand the nature of this O_2_ consumption, we added CN^-^, a tight-binding inhibitor of *aa*_3_-type oxidases, and found that unlike in exponential-phase vegetative cells (Fig 1E), O_2_ consumption in spores was CN^-^-resistant, consistent with previous studies (Powell, 1951; Wilkinson et al., 1977). There is spectroscopic evidence for the presence of an alternative, CN^-^-insensitive oxidase in spores: a cytochrome *bd* oxidase (Hogarth et al., 1977). We confirmed this observation by isolating the inner membrane from *B. megaterium* spores and vegetative cells and measuring the air-oxidised minus dithionite-reduced difference spectrum to reveal the characteristic peaks that arise from different haem groups when they become reduced (Fig 1G). The presence of a 630 nm peak is diagnostic for the *d*-haem of the *bd* oxidase. As cytochrome *bd* is a high affinity terminal oxidase typically used under low O_2_ tension/stress conditions (Friedrich et al., 2022), we were surprised to find it in the spores of *B. megaterium*, an aerobic species cultured in O_2_-replete conditions with vigorous shaking. The genome of both *B. megaterium* and *B. subtilis* encodes two *bd* oxidase enzymes, CydAB (BMQ_5230 & BMQ_5231; BSU_38760 & BSU_38750) and YthAB (BMQ_487 & BMQ_4879; BSU_30710 & BSU_30720) (Winstedt et al., 1998; Winstedt and Von Wachenfeldt, 2000). Running isolated *B. megaterium* membranes, solubilised in the mild detergent DDM, on a BN-PAGE gel and subjecting the bands to mass spectrometry revealed only peptides for YthAB (Fig 1F), consistent with studies of the *B. subtilis* and *B. anthracis* spore membrane proteomes (Chen et al., 2019; Zheng et al., 2016). During aerobic exponential growth of *B. subtilis*, YthAB was absent, found to be of no importance by deletion, and a physiological role for it is yet to be identified (Hederstedt, 2021; Winstedt and Von Wachenfeldt, 2000).

In addition to our discovery of Yth in spores, we could only detect the YumB isoform of the NADH dehydrogenase-quinone reductase, rather than enzymes encoded either by *ndh* (BMQ_2631) or *yutJ* (BMQ_4971). We found proteomic evidence for most of the canonical respiratory enzymes that lack isoforms, which are listed and illustrated in Fig 1F and 1H respectively. The presence of this isoform of NDH-2 and Yth supports the idea that the respiratory apparatus in spores, whilst sharing some enzymes with that of vegetative cells, is attuned for its role in germination.

### Spectral unmixing reveals electron transfer starts before detectable hydration in germinating spores

To understand the kinetics of electron transfer and hydration, we used spectral decomposition to unmix the observed remission spectra containing multiple overlapping peaks into the additive components that correspond to individual haems bound within ETC cytochromes. The haem components were modelled based on the now extensive knowledge of ETCs and the spectra that arise from cytochrome centres (Fig 2A). This analysis allowed us to visualise how individual spectral components changed with respect to each other over time (Fig 2B-F). A smooth baseline was included to account for the large scattering changes that occur as spores rehydrate and undergo morphological changes. To compare our results with literature measurements, we also directly plotted the loss of attenuance (ΔD) at 600 nm that is widely used as a proxy for rehydration in the field with ‘loss of absorbance’ measurements; the underlying physical principle of this measurement is the same.

In the kinetic traces generated for selected spectral components, ΔD at 600 nm, and the OCR (Fig 2G-H), we could distinguish four clusters of events, designated 1-4. Interestingly, reduction of *b* and *c* haem groups (cluster 2) started almost half a minute before ΔD @ 600 nm (cluster 3) and a minute before change in the OCR trace (cluster 4), indicating that electrons are entering the ETC before detectable hydration but are not travelling all the way down the ETC to O_2_. Furthermore, the *aa*_3_-type oxidase F intermediate (cluster 1) accumulated faster than the reduction of *b* and *c* haem groups, which can be attributed to the reduction of Qox to which electrons are delivered directly via the Q-pool, bypassing cytochromes that contain *b* or *c* haems.

It was surprising that the reduction of cytochromes was not immediately followed by an increase in OCR as electrons are normally transferred through ETCs in under a second (Blaza et al., 2014; Trouillard et al., 2011). This suggests that in the ETC of germinating spores, electrons are rapidly passed to the *aa*_3_-type oxidases Qox and Cta but O_2_ reduction does not occur for some reason. To rationalise this observation, we examined the extensively studied catalytic cycle of *aa*_3_-type oxidases (Fig 2I) closely (Wikström et al., 2023). With the delivery of 4 protons and 4 electrons to the BNC, one O_2_ is fully reduced to two H_2_O molecules. Each proton and electron transfer step to the BNC is coupled with the pumping of an additional proton through the membrane, building the ΔΨ or proton-motive force (PMF). Numerous intermediates of haem *a*_3_ coordinating the partially reduced O_2_ are formed during the catalytic cycle at the BNC (termed R ⇾ A ⇾ P_M_ ⇾ F ⇾ O_H_ ⇾ E_H_ ⇾ R), and some of these intermediates give absorbance spectra with peaks at specific wavelengths (Fig 2I). The entire catalytic cycle is reversible except the reduction of O_2_ (A ⇾ P_M_), where the combination of highly oxidised cytochrome *c* and the backpressure from a high PMF can drive the mitochondrial BNC from R to P_M_ (Wikstrom 1981). At a slightly lower PMF, the F intermediate accumulates (Björck and Brzezinski, 2018; Shimada et al., 2023), which is not able to react with O_2_. The earliest recorded events in the *B. megaterium* germination cascade are the release of cations, flowing down a concentration gradient (Racine et al., 1979; Swerdlow et al., 1981). This release is now known to be at least partially mediated by the germinant receptors (Gao et al., 2023); whether this release during germination will build ΔΨ will depend on the counter movement of charge. With the observation of the F intermediate of the *aa*_3_-type oxidases, we now provide evidence that a substantial ΔΨ is very rapidly formed. Establishment of high ΔΨ at the earliest stages of germination offers an explanation for the presence of abundant cytochrome *bd* (Yth): as this enzyme is not proton-pumping, it will not be slowed by ΔΨ to the same extent and allows rapid transfer of electrons to O_2_ as spores germinate.

### Glucose-powered metabolism starts early in germination

In the current model for spore germination, bioenergetic processes resume only after core hydration is complete (Christie and Setlow, 2020; Ghosh et al., 2015; Korza et al., 2023; Setlow and Christie, 2023; Swarge et al., 2020a), incompatible with our data which shows some ETC filling before detectable hydration and O_2_ consumption starting around the time of hydration. Therefore, we wanted to re-evaluate the timeline of metabolic events in the germination cascade to understand how the ETC is being reduced.

We used different concentrations of glucose to germinate spores in the presence and absence of O_2_, using traditional loss of absorbance measurements. As previously observed (Hyatt and Levinson, 1961), the extent and rate of absorbance loss increased with glucose concentration (Fig 3A&B). Unexpectedly, we found that the absence of O_2_ increased the rate of hydration; a likely explanation is that oxidative metabolism could not occur so less glucose was metabolised and more was available to activate GerU. In support of this hypothesis, rehydration with KBr, a non-metabolisable germinant, was unaffected by the absence of O_2_. Outgrowth relied on the presence of both O_2_ and a complete medium (nutrient broth) and was inhibited by ∼60% with 10 mM KCN (Fig 3C-D).

**Figure 3.**
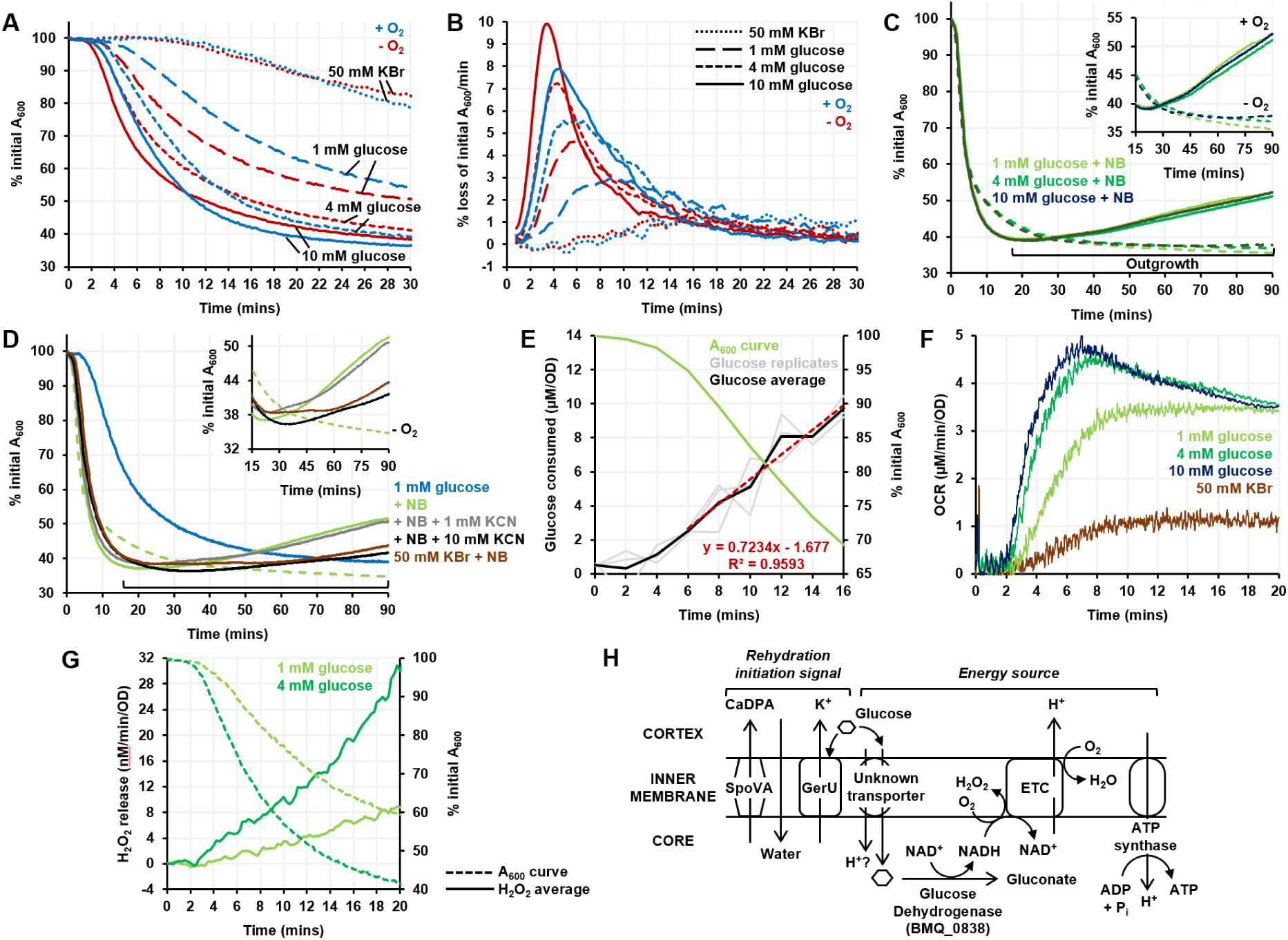
The role of glucose as both a germinant and energy source in germinating spores. (A) Rehydration curves (expressed as % initial absorbance at 600 nm vs. time) for spores when germination is initiated with 50 mM KBr (small dots), 1 mM glucose (long dashes), 4 mM glucose (short dashes) and 10 mM glucose (solid line), under oxic (blue) and anoxic (red) conditions. (B) Rate of spore rehydration (% loss of initial absorbance/min) with 50 mM KBr, 1 mM glucose, 4 mM glucose and 10 mM glucose under oxic and anoxic conditions. (C) Outgrowth curves for spores germinated with nutrient broth and 1 mM glucose (light green), 4 mM glucose (dark green) and 10 mM glucose (teal), under oxic (solid line) and anoxic (dashed line) conditions. (D) Outgrowth curves for spores germinated with 1 mM glucose alone (light blue), 1 mM glucose and nutrient broth (light green), 1 mM glucose and nutrient broth under anaerobiosis (light green dashed), 1 mM glucose, nutrient broth and either 1 mM KCN (grey) or 10 mM KCN (black). (E) Average glucose consumption by spores germinated with 1 mM glucose (black) across replicates (grey) along with the rehydration curve obtained with 1 mM glucose (blue). Glucose was consumed at a rate of 0.72 μM/min (normalised by OD) in the 6-16 minute time period. (F) OCRs measured without the O_2_ delivery system, for rehydrating spores germinated with 1 mM glucose (light green), 4 mM glucose (dark green) and 10 mM glucose (dark blue), and 50 mM KBr (maroon). (G) Rate of H_2_O_2_ release by spores germinated with 1 mM glucose (light green) and 4 mM glucose (dark green). (H) Model for how glucose links the initiation of germination with resumption of bioenergetics in *B. megaterium* spores.

Previous studies came to contradictory conclusions about the timeline of glucose uptake and consumption germination, having found that they start either late (Racine et al., 1979), or early (Maruyama et al., 1980). To establish the timeline in our experimental system we measured the decrease in glucose concentration as spores germinated. Glucose concentration in the buffer medium started decreasing early in germination, demonstrating its rapid consumption (Fig 3E). Additionally, we saw a rapid increase in OCR that was glucose concentration-dependent and started at the same time as rehydration and glucose consumption, i.e. ∼2 minutes after glucose addition (Fig 3F).

In *B. megaterium*, glucose can be catabolised by the glycolytic, pentose-phosphate (PP) and gluconate pathways. The gluconate pathway joins the PP pathway via the intermediate 6P-gluconate (Wushensky et al., 2018). All three pathways become operational in germinating spores of *B. megaterium*, but the gluconate pathway is responsible for more than half of the glucose catabolism in the first 15 mins (Otani et al., 1986). To identify the spore isoform of GDH, we carried out cell fractionation and proteomics. There are three annotated isoforms of GDH in the *B. megaterium* genome: BMQ_0838, BMQ_1051, & BMQ_3371. GDH activity was enriched from the whole soluble fraction of disrupted spores by successive column chromatography steps, and the fraction with the highest enzymatic activity was subjected to LC/MS analysis. BMQ_0838 was found to encode the spore isoform (Fig S1, Table S2, and supplementary .xlsx file). As the gluconate pathway dominates glucose catabolism during rehydration, the majority of glucose taken up is unphosphorylated and its uptake would have to be mediated by a non-PTS (phosphotransferase system) permease. A potential candidate is GlcU where transport is PMF-driven (Castro et al., 2009). *GlcU*, which encodes a non-PTS glucose:H^+^ symporter, is in the same operon as the spore-specifc *gdh* in *B. subtilis* and *B. megaterium*. Both *gdh* and *glcU* are expressed during sporulation in *B. subtilis* (Nicolas et al., 2012). We were unable to find GlcU in our LC/MS analysis but it has previously been identified in the *B. subtilis* spore membrane proteome (Zheng et al., 2016).

An important signature of oxidative metabolism is reactive O_2_ species (ROS) production, where O_2_ adventitiously captures electrons from metabolic enzymes to form unstable superoxide and peroxide species rather than productively forming water at the oxidases. We measured that H_2_O_2_ production started at the same time as glucose metabolism. In the first 30 minutes of germination, ∼0.2 μM and ∼0.7 μM H_2_O_2_ was released by spores germinating with 1 mM and 4 mM glucose respectively (Fig 3G). The H_2_O_2_ measured in germinating spores is an underestimate of the total H_2_O_2_ production as the majority would be neutralised by catalase, but the proportional increase in H_2_O_2_ released supports the idea that in germinating spores oxidative metabolism is ongoing from the earliest stages of germination.

Finally, our hypothesis for glucose-powered oxidative metabolism during germination relies on the NADH produced being able to rapidly reduce the ETC. Kinetic assays showed that spore membranes oxidised NADH nearly twice as fast as membranes from vegetative cells (0.26 versus 0.15 μmol min^-1^ mg^-1^ protein), demonstrating that the spore ETC is ready to accept electrons from NADH. Spore membranes were also more CN^-^-resistant, consistent with the function of CN^-^-resistant Yth [Table S1].

### The *ythAB*-encoded cytochrome *bd* oxidase drives ETC function in germinating spores

We have established that the spore ETC is rapidly reduced early in germination, powered by glucose metabolism. We now turn to the role of the spore-specific *bd* oxidase, Yth. Traditionally considered O_2_-scavenging enzymes required for growth in microaerobic conditions, the role of a *bd* oxidase in spores cultivated and germinated in O_2_-replete conditions was mysterious. Given that Qox and Cta are operating in distinct and slower catalytic regimes compared to vegetative cells, we speculated that Yth could be acting to relieve reductive pressure.

To test the hypothesis that Yth drives ETC function when the haem-copper oxidases are hindered, *ythA*, the haem-containing catalytic subunit of the heterodimeric cytochrome *bd* oxidase, was deleted in *B. megaterium* spores (Fig 4A). Deletion of *ythA* was confirmed by remission spectroscopy on membranes isolated from Δ*ythA* spores (Fig 4B). Compared to WT, Δ*ythA* spores underwent slower rehydration when germinated with 10 mM glucose (Fig 4C) but achieved a comparable OCR (Fig 4D). However, unlike in WT spores (Fig 4D), O_2_ consumption in germinating Δ*ythA* spores was fully abolished by 1 mM KCN (Fig 4E), consistent with the deletion of a CN^-^-resistant *bd* oxidase without induction of Cyd, the other *bd* oxidase in the genome. Difference spectra in Fig 4F-H show that compared to WT, Δ*ythA* spores had less of the 580 nm F species, and the small feature at 600 nm became stronger when KCN inhibited O_2_ consumption and reduced the haem *a* centres further. This change suggests that in the absence of Yth, the *aa*_3_-type oxidases were pushed to operate faster and some of the 600 nm intermediate accumulated, as it does in vegetative cells.

**Figure 4.**
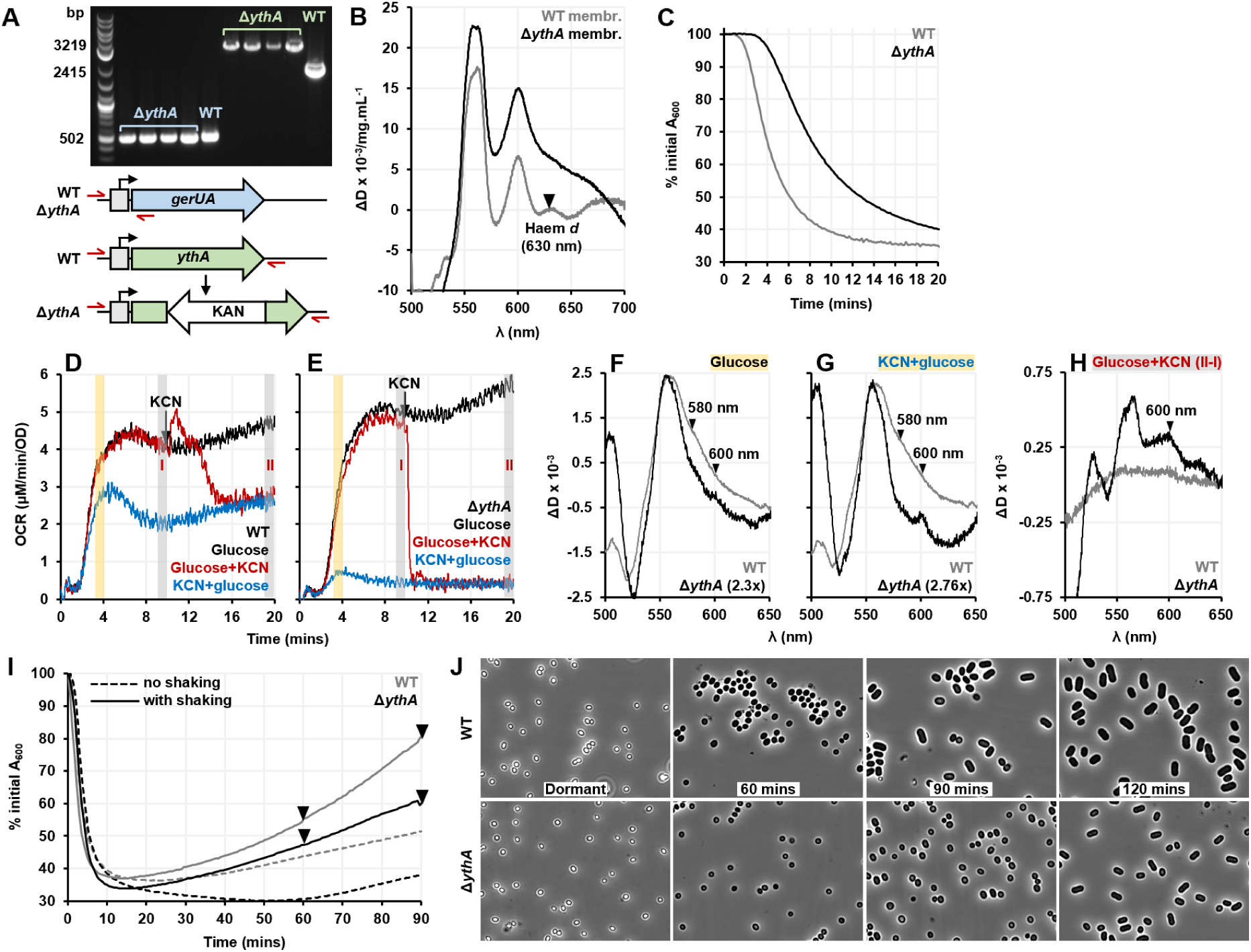
Effect of the *ythAB*-encoded cytochrome *bd* oxidase deletion on *B. megaterium* spore germination. (A) The WT and Δ*ythA* genotypes confirmed by PCR. Both strains have the plasmid-borne glucose receptor GerU but Δ*ythA* is an insertional deletion mutant of *ythA*. (B) The deletion of cytochrome *bd* oxidase confirmed by haem remission spectroscopy. The regression-subtracted dithionite-reduced minus air-oxidised spectra of isolated membranes from dormant WT (grey) and Δ*ythA* (black) spores. (C) Rehydration curves obtained for the WT (grey) and Δ*ythA* (black) spores germinated with 10 mM glucose. (D) OCRs measured for WT and (E) Δ*ythA* spores germinated with 10 mM glucose (black), when 1 mM KCN was added either before glucose (blue) or 10 mins after glucose (red). (F) Difference spectra generated for WT (grey) and Δ*ythA* (black) spores as described in Fig 1B. (F) and (G) show the T_p_=4 min spectra (T_p_ minus pre-glucose) indicated by the yellow shaded region in (D) & (E), when spores were germinated either with glucose alone (F) or in the presence of KCN (G). (H) shows the difference spectra generated by subtracting I from II, indicated by grey shaded regions in (D and (E), when KCN was added 10 mins after glucose. (I) Outgrowth curves for WT (grey) and Δ*ythA* (black) spores germinated with 10 mM glucose and nutrient broth, when the 96-well plate was shaken (solid line) or not (dashed line) during the experiment. (J) Phase-contrast micrographs of WT (top panel) and Δ*ythA* (bottom panel) spores taken at the time points indicated by arrowheads during the outgrowth experiment (with shaking) shown in (I).

Finally, phase-contrast imaging was used to follow germination of WT and Δ*ythA* spores. Before imaging, spores were germinated in a 96-well plate (Fig 4I) and transferred to a glass slide for imaging with phase-contrast microscopy. For Δ*ythA* spores, outgrowth was much slower when the 96-well plate was not shaken, as the oxygenation was insufficient and the high-affinity cytochrome *bd* oxidase was also absent. With shaking, both WT and Δ*ythA* spores were able to outgrow better but Δ*ythA* cells still lagged behind WT cells as seen in the outgrowth curves and the accompanying phase contrast micrographs (Fig 4J).

### Demonstrating that similar principles underlie *Bacillus subtilis* spore germination

Next, we attempted to reproduce these findings in WT and Δ*ythA B. subtilis*. As observed in *B. megaterium*, the Δ*ythA B. subtilis* spores were slower to rehydrate than WT when germinated with 10 mM alanine (Fig 5A), directly linking Yth and bioenergetic processes to hydration. However, when germination was initiated with alanine in the absence of glucose, neither strain was able to achieve an appreciable OCR (Fig 5B & C). The addition of equimolar glucose along with alanine led to a considerably higher OCR and more pronounced spectral changes in both WT and Δ*ythA B. subtilis* spores, supporting the idea that while interrelated, germinant recognition and resumption of bioenergetics can progress independently. As expected, O_2_ consumption was abolished by KCN in Δ*ythA B. subtilis* spores as flux is forced to the CN^-^-sensitive *aa*_3_-type oxidases (Fig 5C).

**Figure 5.**
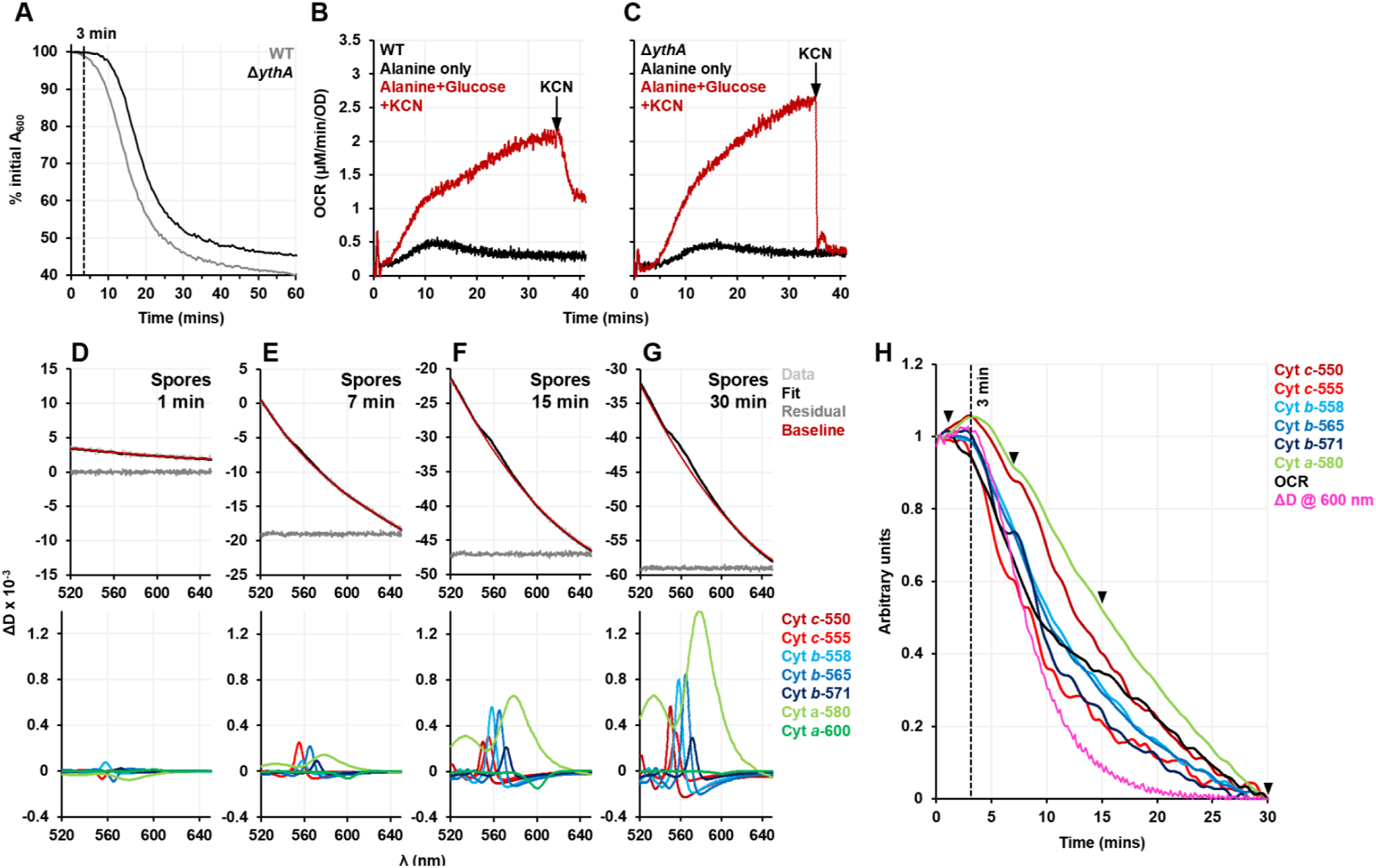
Translation of findings from *B. megaterium* in the model spore-former *B. subtilis*. (A) Rehydration curves measured for the WT (grey) and Δ*ythA* (black) spores germinated with 10 mM alanine. OCRs measured for (B) WT and (C) Δ*ythA* spores germinated with 10 mM alanine alone (black) and 10 mM alanine+glucose (red). In the latter experiments 1 mM KCN was also added at T_p_=35 mins. (D)-(G) are spectra of spores germinated with 10 mM alanine+glucose. The top panel shows the raw data, the fit imposed, the residuals (suitably offset from 0 to be comparable), and the sum of the baseline components being fitted at the indicated time points. The bottom panel shows the fitting of the haem components. (H) Savitzky-Golay smoothed and normalised kinetic traces (fitting versus time) of haem components, change in attenuance (ΔD) at 600 nm and OCR for spores germinated with 10 mM alanine+glucose. Arrowheads indicate the time points 1, 7, 15, and 30 mins for which the decomposition analysis is demonstrated in (D)-(G).

**Figure 6.**
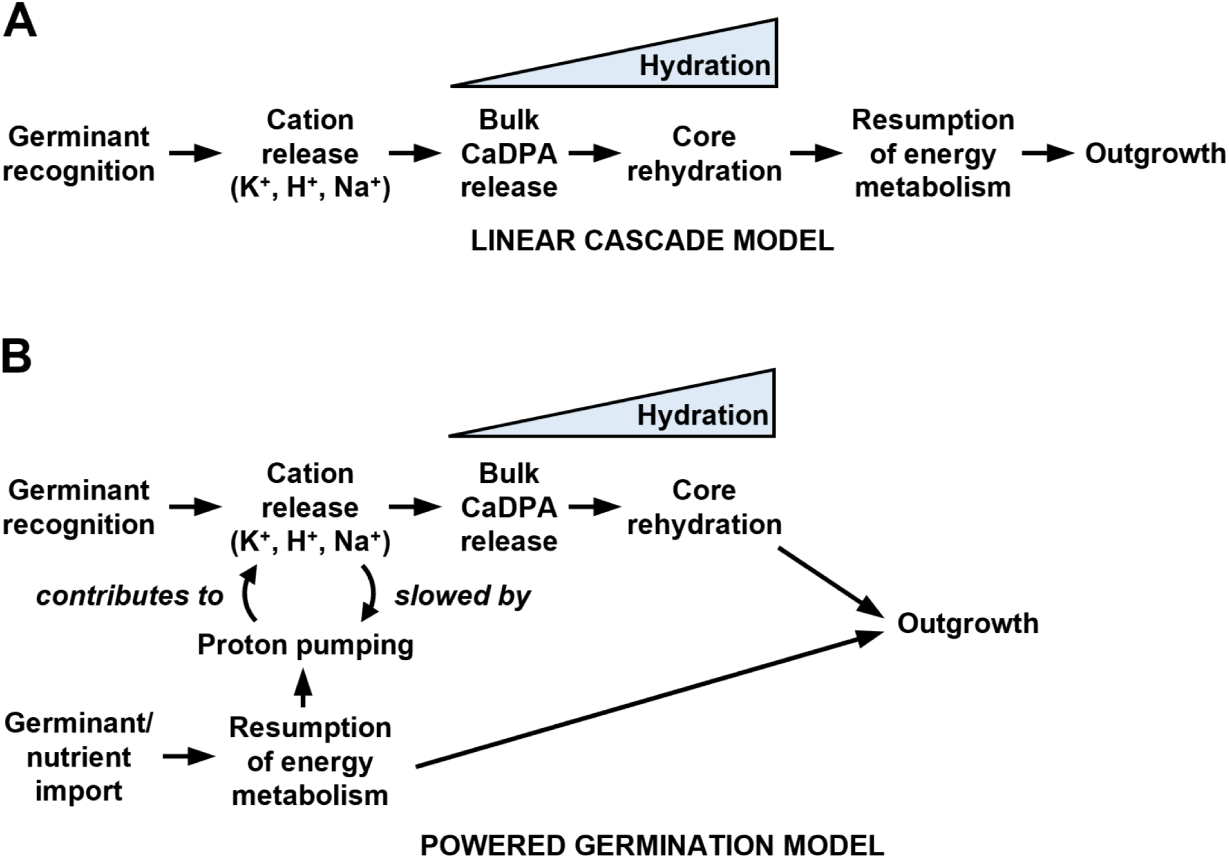
The linear cascade and powered germination models of *Bacillus* spore germination. (A) The linear cascade model of spore germination in which energy metabolism is resumed only after spore rehydration is completed. (B) The powered germination model based on the findings in this work. A nutrient germinant is recognised by the cognate germinant receptor which induces cation release and it can simultaneously be imported into the spore where its catabolism powers the resumption of electron transport and proton pumping.

Germinating *B. subtilis* spores presented different remission spectra compared to *B. megaterium*. In *B. subtilis*, the changes in attenuance were slower and approximately half that measured for *B. megaterium* spores at the same spore density in the bioenergetic chamber (Fig 5D-G). In *B. megaterium*, the change in baseline presented mostly as an even loss of attenuance across the spectral range, with a minor element of a quadratic function creating a ‘U’, modelled in Fig 2A as γ = (x-585)^2^. In *B. subtilis* spores, there was a loss of attenuance across the spectrum, but only a half of the quadratic ‘U’ was observed, modelled with α = 1/x^2^. We lack a firm explanation for this behaviour but note that while spores of the two species have similar volumes (0.47±0.07 μm^3^ for *B. subtilis*; 0.51±0.11 μm^3^ for *B. megaterium*), they are distinct in their aspect ratios, with *B. subtilis* being more elongated (1.73 ± 0.17 for *B. subtilis*; 1.26 ± 0.17 for *B. megaterium*) (Xu Zhou et al., 2017). The wavelength of the light used in the chamber and the dimensions of the spores are of the same order (∼1 μm) so the scattering processes will depend on the morphology of the spores (Harding, 1997). Despite the differences, the unmixing model was sufficiently flexible to work well for *B. subtilis* spores, providing good solutions judged by low residuals (Fig 5D-G, top panels). In *B. subtillis*, we found that rehydration, haem reduction, and O_2_ consumption all start simultaneously. Intriguingly, compared to *B. megaterium*, OCR in *B. subtilis* started increasing earlier concomitantly with haem reduction, and the F-intermediate which is indicative of ΔΨ, built much slower (Fig 5H). Therefore, either ΔΨ is lower in germinating *B. subtilis* spores, or the balance of the two *aa*_3_-type oxidases favours Qox, which experiences a higher Δ*E* (the thermodynamic ‘driving force’ across the enzyme) than Cta and is better ‘powered’ to overcome the backpressure of ΔΨ. This increased driving force is because the Qox pathway passes electrons from menaquinone (*E*_m_ = −74 mV) to O_2_ (*E*_m_ = +840 mV), giving a Δ*E*_m_ of 914 mV, whilst Cta passes electrons from cytochrome *c* (*E*_m_ = +220 mV), giving a Δ*E*_m_ of 620 mV (Nicholls and Ferguson, Stuart J, 2002).

Finally, for Δ*ythA B. subtilis* spores, outgrowth progressed as in WT spores, and shaking did not make any meaningful difference (Fig S2). For outgrowth to occur, germinating spores require a complete medium like nutrient broth, which may provide the Δ*ythA B. subtilis* spores other metabolisable amino acid energy sources that decrease dependence on oxidative metabolism. *B. megaterium* spores must also conserve more energy to build a bigger vegetative cell than *B. subtilis* spores, which would make *B. megaterium* spores particularly sensitive to perturbations of bioenergetic pathways, especially when they are already energetically depleted as they exit dormancy.

## Discussion

The data in this work leads us to propose the following model for bacterial spore germination in *B. megaterium* and *B*. *subtilis*, which we call the powered germination model. Germinants can act both as a signal and as an energy source that is immediately exploited; cytoplasmic metabolism can occur in the gel-like core of a yet-to-be-hydrated spore. In *B. megaterium*, glucose recognition by GerU causes cation release; simultaneously, glucose is imported into the spores where it is metabolised, passing electrons to the ETC. In *B. subtilis*, the role of glucose is predominantly to act as an energy source, and alanine is recognised by the GerA receptor (Prasad et al., 1972). Cation efflux and glucose-powered proton pumping causes a substantial ΔΨ to be rapidly established, which we detect as it causes the F-intermediate in the catalytic cycle of *aa*_3_-type oxidases to accumulate. Our approach cannot inform on the presence or absence of ΔΨ before the *aa*_3_-type oxidases are reduced but we can observe that ΔΨ is present from the earliest stages of germination, certainly before hydration in *B. megaterium*, where most likely ΔΨ is built by the Ger-mediated efflux of cations. The substantial ΔΨ requires that these spores possess Yth; as Yth does not pump protons it will be insensitive to ΔΨ, allowing the oxidative metabolism of glucose even when *aa*_3_-type oxidases are slowed down by ΔΨ. The receptor-mediated signalling cascade and bioenergetic processes are semi-independent as demonstrated by the deletion of *yth* slowing hydration.

Our model is a substantial departure from the current consensus, which holds that germinating spores cannot support metabolic processes until cortex degradation and core rehydration is complete (Christie and Setlow, 2020; Ghosh et al., 2015; Korza et al., 2023; Setlow and Christie, 2023; Swarge et al., 2020a). A central tenet in the field is that during the early stages of germination, most of the molecular processes are running spontaneously downhill, such as cation/CaDPA release, water ingress, and proteolysis. Our findings allow for powered processes such as active transport to occur early in germination. Bioenergetics are relevant to areas of spore biology where the energetically-depleted nature of spores is invoked as a limiting factor, e.g. arguments regarding protein synthesis occurring early in germination (Setlow and Christie, 2020; Zhou et al., 2022).

Orthogonal approaches support our powered germination model. Recently, an endogenous luciferase-luciferin system was established in *B. subtilis* to study the dynamics of energy metabolism, where the bioluminescence was directly linked to the redox potential of individual germinating spores;. the bioluminescence signal was synchronised with phase transition (Frentz and Dworkin, 2020). In *B. atrophaeus*, O_2_ was found to slow germination, consistent with Figure 3, but was essential for subsequent colony formation (Wu and Chang, 2022).

There is a substantial body of work concluding that bioenergetic processes become operational only later in the germination process. ATP accumulation has been measured on extracts from germinating spores (Dills and Vary, 1978; Korza et al., 2023; Setlow and Kornberg, 1970). These studies found that ATP only accumulates 1-2 minutes after hydration starts, concluding that ATP synthesis is not required for the initiation of germination. However, steady-state ATP concentration can be low if ATP production is high but consumption is even higher, particularly if starting from a point of low ATP, a situation anticipated by the original authors (Scott and Ellar, 1978). But in spores exiting dormancy, the concentration of metabolic intermediates for reactions powered through ATP hydrolysis will be far removed from those found within a vegetative cell, so in many cases the phosphorylation potential required to drive reactions will be lower as metabolic pathways are empty. Additionally, Yth has a role in ensuring rapid oxidation of the ETC, which will impose an oxidative ‘pull’ on central metabolism.

Our model requires that metabolites are sufficiently mobile for reactions to occur. Compared to vegetative cells, molecular diffusion is decreased in spores but careful biophysical experiments have demonstrated that some molecular motion still occurs. While proteins are rotationally immobilised, water remains highly mobile, thus the description of the core as a ‘gel’ rather than a ‘glass’ (Kaieda et al., 2013; Sunde et al., 2009). It is plausible that small molecules, such as glucose, will have sufficient mobility to diffuse between the immobilised metabolic enzymes. A similar principle applies to lipids immobilised in the spore cytoplasmic membrane (Cowan et al., 2004).

There has been renewed interest in ion movements during spore germination, particularly cation efflux (Gao et al., 2023; Kikuchi et al., 2022). While these have been seen as signalling events, such charge movements are also a fundamental form of energy conservation. There are precedents for this dual role, for example, it has been observed that the ATP synthase inhibitor DCCD delays the release of CaDPA and the re-uptake of K^+^ ions in *B. megaterium* spores (Swerdlow, Setlow and Setlow, 1981). Proton pumping by the *aa*_3_-type oxidases will contribute to both the PMF to power ATP synthesis and the imbalance of charge that the Ger receptors create. In *B. megaterium*, there is little O_2_ consumption until cytochrome reduction is well underway, so proton pumping will contribute less to the Ger-mediated cation efflux, but in *B. subtilis*, O_2_ consumption starts relatively earlier alongside cytochrome reduction, so proton pumping may contribute directly to the downstream activation of SpoVAD channels which release CaDPA in response to the germinant signal (Gao et al. 2023).

Depots of the glycolytic intermediate 3-PGA (3-phosphoglyceric acid) are thought to be the major endogenous energy reserve that spores rely on during germination as substrate-level phosphorylation of 3-PGA-derived PEP generates ATP. Spores germinated in nutrient-free media with KBr (Setlow and Kornberg, 1970) or CaDPA (Korza et al., 2023b) are reliant on this pathway. Indeed, in the inverse process by which dormany is established, recent modelling suggests that the forespore does generate ATP by substrate-level phosphorylation of glycolytic intermediates provided by the mother cell (Tibocha-Bonilla et al., 2025). However, spores germinated with physiological germinants will have a substantial exogenous energy supply to be exploited for higher ATP yields by oxidative phosphorylation. In *B. subtilis*, alanine alone leads to rapid hydration but relatively low O_2_ consumption and meagre cytochrome reduction, but Δ*yth* still slows alanine-initiated hydration because alanine transamination/oxidation and gluconeogenesis also subsequently lead to ETC reduction (Prasad, 1974). Thus, the potency of a germinant can be partially linked to the capacity a germinating spore has to directly use it, e.g. glucose is rapidly catabolised by the gluconate pathway in *B. megaterium*, to generate reducing power. Alanine is ubiquitously recognised as a germinant by Bacillus and Clostridial spores and even some fungi, suggesting it is a common ancestral germinant (Ijadpanahsaravi et al., 2024; Ross and Abel-Santos, 2010). We speculate that alanine is a reliable environmental signal for many of these spores but might not be as readily metabolisable by existing pathways, which in *B. subtilis* also favour the co-germinant glucose as the source of reducing power (Prasad et al., 1972).

The cytochrome *bd* oxidases are increasingly recognised as being central to bacterial physiology (Friedrich et al., 2022). Well-established as O_2_ scavenging enzymes (Miller and Gennis, 1983), the cytochrome *bd* oxidases have emerging roles as reductive pressure ‘relief valves’. Starting with the observation that a deletion of the mycobacterial *bd* oxidase, Cyd, increases sensitivity to the ATP-synthase inhibitor bedaquiline (Berney et al., 2014), we recently used the bioenergetic chamber to show that even for mycobacteria cultured in plentiful O_2_, the *bd* oxidase allows the relief of reductive pressure created by bedaquiline (Harrison et al 2024). Our observations here support Yth having a very similar role in germinating spores. Having two divergent physiological contexts where the *bd* oxidases function as a pressure relief valve emphasises this new role.

While the connection between bioenergetics and germination was explored early in the development of the field, technical limitations meant it was overlooked and discounted. Our measurements on germinating spores bypass the caveats created by quench and measure approaches, and allow us to directly study spores responding to germinants. Here we have focussed on two model species, *B. megaterium* and *B. subtilis*, but we anticipate that similar principles will apply to many other spore-formers.

## Acknowledgements

This work is supported by a UKRI Future Leader Fellowship to JNB (MR/T040742/1) and a BBSRC sLoL award (BB/X003035/1). PG was supported by a York Overseas Scholarship and the Department of Chemistry, University of York. RCW was supported with a BBSRC White Rose PhD Studentship (2434192). ECW and BLH carried out the work as part of their Masters degrees at the University of York, in Biochemistry and Chemistry, respectively. We thank Jules Borgia and Roxy Osinska in the YSBL wet labs, Dr Adam Dowle and Dr Chris Taylor in the the Centre for Excellence in Mass Spectrometry, and Dr Andrew Leech in the Molecular Interactions Laboratory for their invaluable assistance. The Orbitrap Fusion in the Centre of Excellence in Mass Spectrometry was funded by Science City York (Yorkshire Forward) and EPSRC (EP/K039660/1, EP/M028127/1).

## CRediT (Contributor Roles Taxonomy)

|  | PG | RC | RCW | ECW | BH | GC | RJS | JNB |
| --- | --- | --- | --- | --- | --- | --- | --- | --- |
| Conceptualization | ✓ |  |  |  |  | ✓ | ✓ | ✓ |
| Methodology | ✓ | ✓ | ✓ |  |  | ✓ | ✓ | ✓ |
| Software |  |  | ✓ |  |  |  | ✓ |  |
| Validation | ✓ | ✓ | ✓ |  |  | ✓ | ✓ | ✓ |
| Formal analysis | ✓ | ✓ | ✓ |  |  |  |  | ✓ |
| Investigation | ✓ | ✓ | ✓ | ✓ | ✓ |  |  |  |
| Resources |  |  |  |  |  | ✓ | ✓ | ✓ |
| Data Curation | ✓ | ✓ |  |  |  |  |  |  |
| Writing - original draft | ✓ |  |  |  |  |  |  | ✓ |
| Writing - review and editing | ✓ | ✓ | ✓ | ✓ | ✓ | ✓ | ✓ | ✓ |
| Visualisation | ✓ |  |  |  |  |  |  |  |
| Supervision |  |  |  |  |  | ✓ | ✓ | ✓ |
| Project administration |  |  |  |  |  | ✓ | ✓ | ✓ |
| Funding acquisition |  |  |  |  |  | ✓ | ✓ | ✓ |

## Conflict of interest statement

RJS is the founder of Cellspex Ltd, which is commercialising the remission spectroscopy technology used in this work by developing the ‘Iberius’ bioenergetic chamber. None of the other authors have a conflict of interest.

## Supplementary results

**Table S1.** NADH oxidation rates and comparable % vesicle inversion obtained across biological replicated for spore membranes isolated using the optimised spore membrane isolation protocol.

| **Membranes** | **NADH oxidation rate ($\mu\text{mol}/\text{min}/\text{mg}$)** | **NADH oxidation rate in presence of 1 mM KCN ($\mu\text{mol}/\text{min}/\text{mg}$)** | **Fraction NADH oxidation lost in the presence of 1 mM KCN (%)** |
| --- | --- | --- | --- |
| Vegetative Cells | $0.15 \pm 0.01$ | $0.074 \pm 0.002$ | 51 |
| Spore preparation 1 | $0.23 \pm 0.01$ | $0.20 \pm 0.01$ | 13 |
| Spore preparation 2 | $0.22 \pm 0.01$ | $0.17 \pm 0.01$ | 23 |
| Spore preparation 3 | $0.32 \pm 0.01$ | $0.25 \pm 0.01$ | 22 |

**Fig S1.**
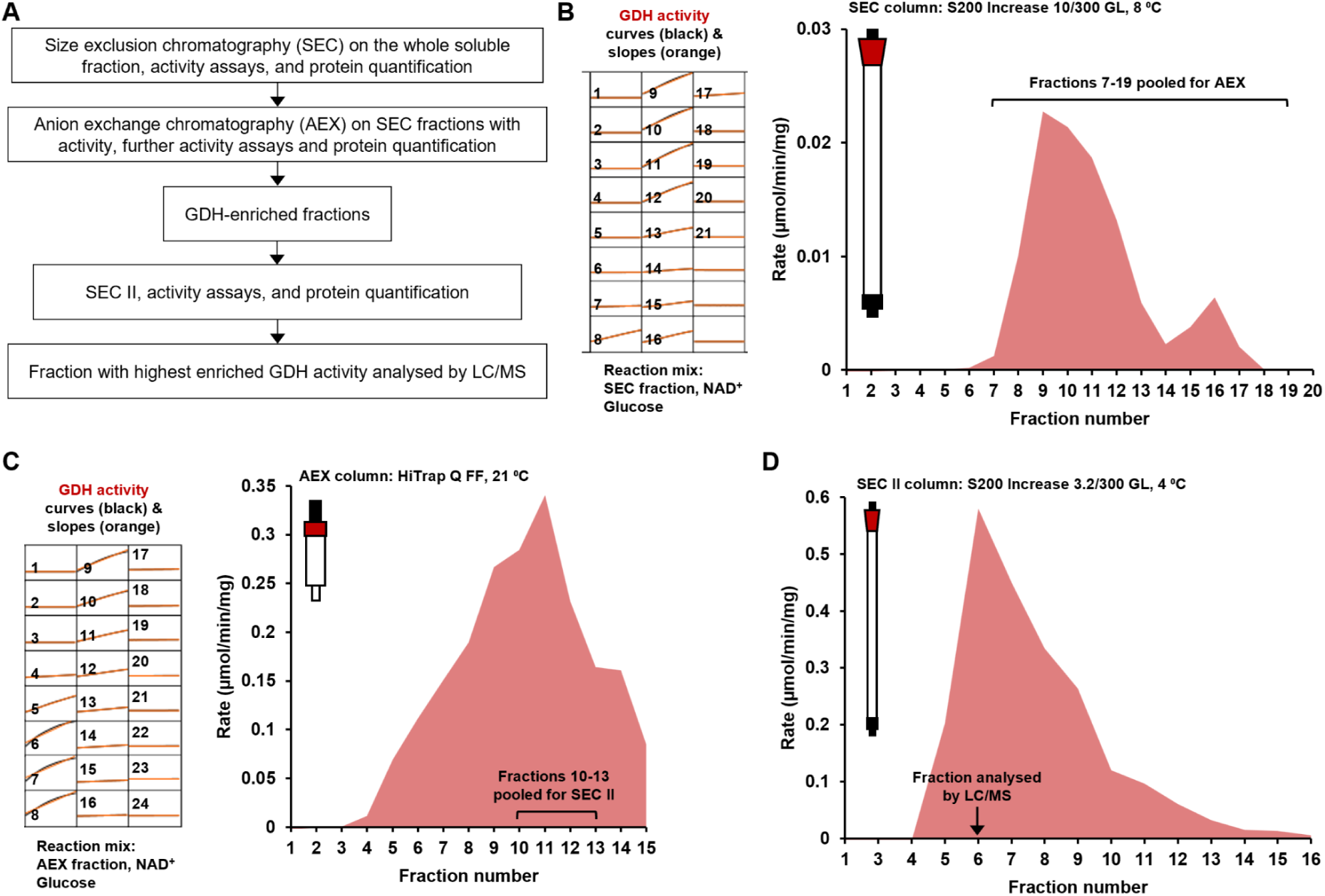
Enrichment of GDH activity from the whole spore soluble fraction. (A) A flowchart summarising the experimental strategy to enrich GDH activity. (B) Left panel shows curves (black) from the spectrophotometric GDH activity assay carried out for each of the SEC elution fractions, the slopes (orange) calculated by the plate reader software, and the components of the reaction. GDH activity would reduce NAD^+^ to NADH, increasing the A_340-380_. The graph on the right shows the rate of GDH activity measured for each fraction numbered 1-21. Based on this, SEC elution fractions 7-19 were pooled and subjected to AEX. (C) Same as (B) but with AEX elution fractions. ‘GDH-enriched’ AEX elution fractions 10-13 were pooled and subjected to another SEC run. (D) After the second SEC, specific GDH activity was enriched further. Fraction GDH-6 was saved for LC/MS analysis.

**Table S2.** Most abundant proteins containing the word “glucose” in the GDH-enriched ‘GDH-6’ fraction identified by LC/MS.

| **Name of protein from UniProt** | **BMQ code** | **Unique peptides** | **KEGG/GenBank annotation** |
| --- | --- | --- | --- |
| Glucose-6 phosphate isomerase | BMQ_4937 | 23 |  |
| Glucose 1-dehydrogenase (NAD(P)<sup>(+)</sup>) | BMQ_0838 | 8 | Glucose 1-dehydrogenase (gdh) |
| Glucose-6-phosphate 1-dehydrogenase | BMQ_5210 | 20 | - |
| Glucose 1-dehydrogenase (NAD(P)<sup>(+)</sup>) | BMQ_2939 | 5 | short-chain dehydrogenase/reductase |
| PTS system, glucose-specific IIBC component | BMQ_1302 | 24 | - |
| Glucose 1-dehydrogenase (NAD(P)<sup>(+)</sup>) | BMQ_2333 | 2 | - |
| Glucose-6-phosphate 1-dehydrogenase | BMQ_1958 | 7 | - |
| Sugar phosphotransferase system, glucose subfamily, IIA component | BMQ_4019 | 3 | - |
| Glucose 1-dehydrogenase (NAD(P)<sup>(+)</sup>) | BMQ_2208 | 3 | 3-hydroxybutyrate dehydrogenase |
| UTP--glucose-1-phosphate uridylyltransferase | BMQ_5130 | 7 | - |
| Glucose 1-dehydrogenase (NAD(P)<sup>(+)</sup>) | BMQ_1269 | 4 | oxidoreductase, short chain dehydrogenase/reductase family protein |

**Fig S2.**
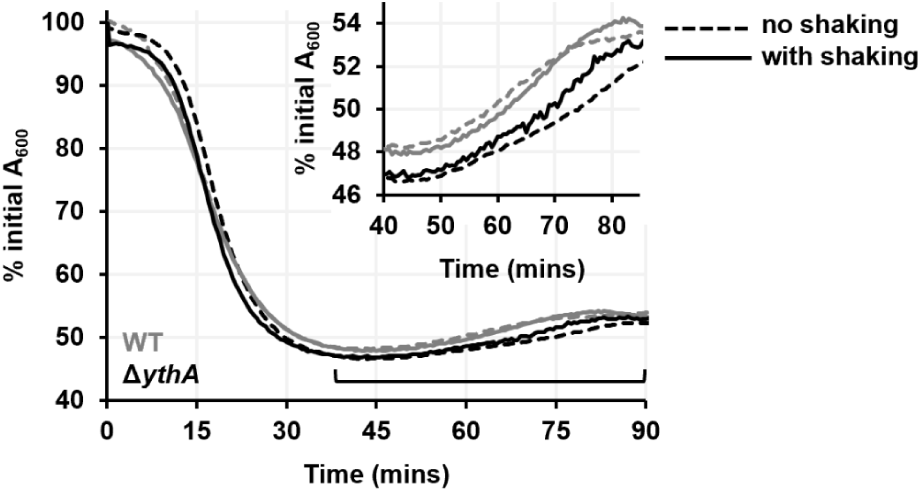
Outgrowth curves for WT (grey) and Δ*ythA* (black) spores germinated with 10 mM alanine and nutrient broth, when the 96-well plate was shaken (solid line) or not (dashed line) during the experiment. Inset shows only the indicated outgrowth phase (40-90 mins) for clarity.

### Methods

#### Bacterial strains and culture conditions

All strains used in this work are listed in the table below along with their genotypic/phenotypic description. The Δ*ythA* mutants were isogenic with the wild-type *B. megaterium* QM B1551 and *B. subtilis* 168 strains. All *Bacillus* strains were routinely cultured in LB medium supplemented with appropriate antibiotics when warranted (erythromycin at 0.5 or 1 μg/mL; lincomycin at 12.5 or 25 μg/mL; kanamycin at 5 or 10 μg/mL; tetracycline at 12.5 μg/mL), and incubated at 30 °C for *B. megaterium* or 37 °C for *B. subtilis*. Plasmids to generate the Δ*ythA B. megaterium* mutant were propagated and isolated from NEB Turbo *Escherichia coli* (New England Biolabs, UK) cultured at 37 °C in LB medium supplemented with carbenicillin (100 μg/mL). The *B. subtilis* WT and Δ*ythA* spores were obtained from the Bacillus Genetic Stock Center library [Koo et al., 2017].

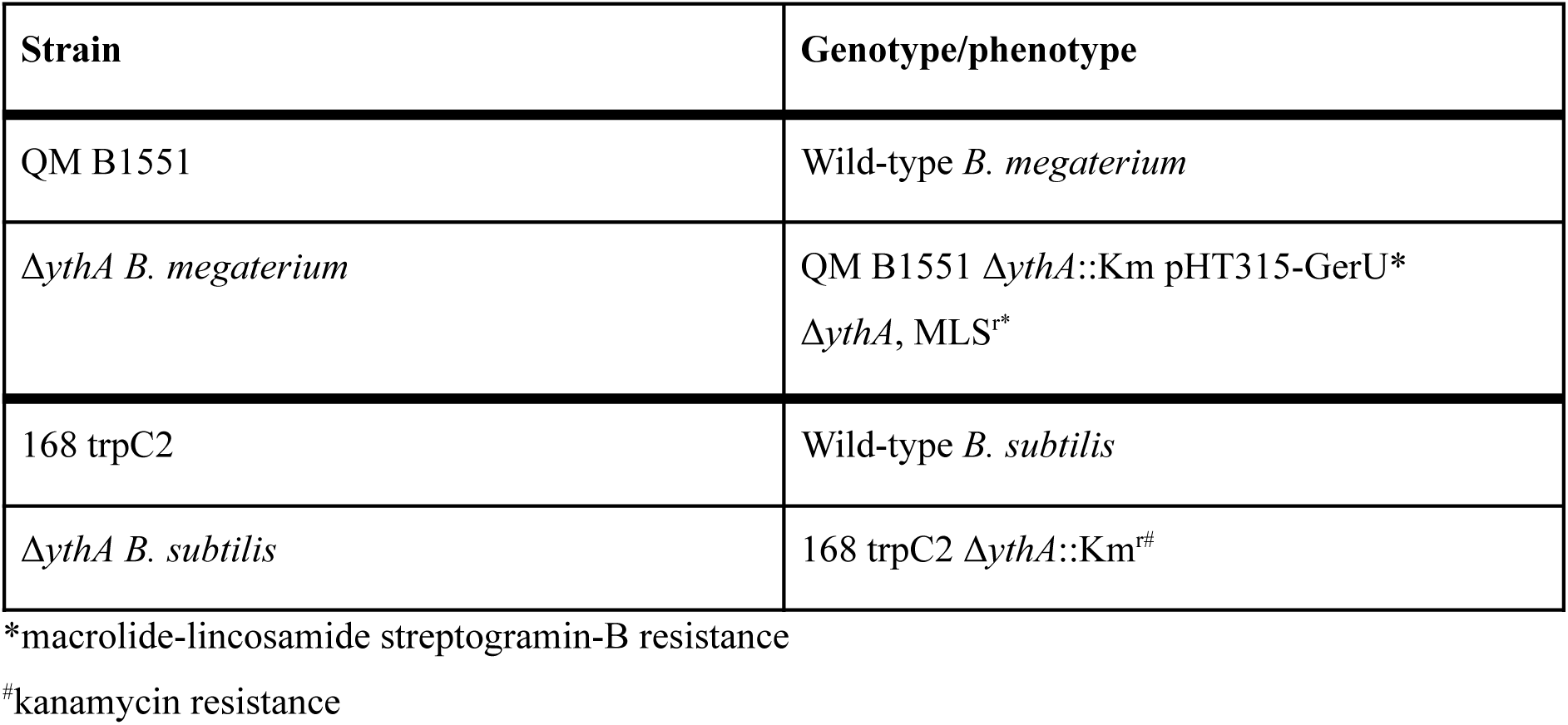

#### Mutant strain construction

An insertion-deletion in ythA (BMQ_4878) of *B. megaterium* QM B1551 was produced by allelic exchange. The plasmid used was assembled from the following, all containing 20 bp overlaps: upstream (positions 4,684,943 to 4,685,552; KEGG *B. megaterium* QM B1551 genome) and downstream (positions 4,685,727 to 4,686,337; KEGG *B. megaterium* QM B1551 genome) regions of BMQ_4878, kanamycin (Km) resistance cassette from pDG792 (Guérout-Fleury et al., 1995), and backbone fragments from pUCTV2 to obtain a temperature sensitive origin of replication and a tetracycline (Tet) resistance cassette (Wittchen and Meinhardt, 1995). PCR products were electrophoresed on agarose gels, extracted under blue light illumination, and purified (QIAquick Gel Extraction Kit, Qiagen, UK). Resultant fragments were combined using Klenow Assembly Method [Bailey and Mohamed, 2018] at 37 °C for 45 mins, with 2 μL of the subsequent product used to transform Turbo *E. coli* (NEB, UK) to carbenicillin resistance. Transformant *E. coli* was harvested and the purified plasmid (QIAprep Spin Miniprep Kit, Qiagen, UK) was verified by sequencing.

Plasmid pUCTV2-ΔBMQ_4878::Km was introduced into *B. megaterium* QM B1551 by polyethylene glycol (PEG)-mediated protoplast transformation, and recovered on RHAF agar containing 5 μg/mL Km at 30 °C overnight [Christie and Lowe, 2008]. Transformant colonies were dotted in a grid-like fashion on LB agar containing 5 μg/mL Km at the non-permissive temperature of 42 °C overnight, to enable integration of the plasmid at the cloned locus via homologous recombination. Sub-culture of the resultant single crossover *B. megaterium* was continued at 42°C on LB agar lacking antibiotics until the desired double-crossover colony was isolated. Successful double-crossover resulted from the excision of pUCTV2 plasmid (Tet^s^) and disruption of BMQ_4878 because of ΔBMQ_4878::Km integration (Km^r^). Colony PCR confirmed the double-crossover mutant but also revealed loss of the native pBM700 plasmid carrying the GerU germinant receptor. To restore the GerU-mediated germination response to glucose, the episomal pHT315-GerU* plasmid [Christie, Lazarevska and Lowe, 2008] was introduced by PEG-mediated protoplast transformation into *B. megaterium* ΔBMQ_4878::Km on RHAF (1 μg/mL erythromycin + 25 μg/mL lincomycin) at 30 °C overnight, and spores were made directly from the resultant transformants.

#### Spore cultivation and purification

*B. megaterium* spores were prepared by nutrient exhaustion in 500 mL of supplemented nutrient broth (SNB) medium [Shay and Vary, 1978; English and Vary, 1986] in baffled flasks shaken at 225 rpm, 30 °C, for 72 hours. For the Δ*ythA* mutant, the SNB medium was supplemented with 0.5 μg/mL erythromycin to maintain the pHT315-GerU* plasmid. The spores were washed 5-6 times (4K RCF, 4 °C, 10 minutes) with ice-cold sterile water until 99% purity was achieved, as confirmed by phase contrast microscopy. The pellet was resuspended in 5 mL sterile water and stored on ice in the cold room at 4 ⁰C.

Washed spores were subjected to further purification using a Histodenz density gradient (Setlow, 2019). 250 μL of the washed spore suspension was centrifuged and the pellet resuspended in 200 μL 20% Histodenz (Sigma-Aldrich), which was layered on top of 1.2 mL 60% Histodenz in a 2 mL microcentrifuge tube. This was centrifuged at 16K RCF for 15 mins at 4 ⁰C, after which the supernatant containing vegetative debris was carefully removed. The purified spore pellet was resuspended, transferred to a fresh tube, and washed 3-4 times with ultrapure water at 4K RCF, 4 ⁰C to remove residual Histodenz after which the purified spores were resuspended in 250 μL ultrapure water. The OD (optical density at 600 nm) of such Histodenz-purified spore suspensions varied between 200-400. Purified spores were always stored on ice and all experiments were performed within 10 days of spore harvest.

*B. subtilis* spores were also prepared by nutrient exhaustion in 2x Schaeffer’s-glucose (2xSG) agar medium (Leighton and Doi, 1971), incubated at 37 °C for 72 hours. For the Δ*ythA* mutant, the 2xSG medium was supplemented with 5 μg/mL kanamycin. The spores were scraped off the plates, washed 5-6 times (18K RCF, 4 °C, 5 minutes), and resuspended in 3 mL sterile water for storage on ice in the cold room at 4 °C. They were later purified using a Histodenz density gradient in the same way as described for *B. megaterium* spores, except that 50% Histodenz was used instead of 60% Histodenz, and the Histodenz-purified spores were stored at an OD of 150.

#### Spore and cell membrane isolation and protein quantification

Membranes were isolated by disrupting spores (from 2 L culture) that were washed thoroughly but not purified further using gradient centrifugation. The washed spores were first treated with 0.1 M NaCl, 0.1 M NaOH, 0.1 M DTT and 0.5% SDS at 37 °C for 1 hour to chemically remove the protein coat and the outer membrane. The de-coated spores were washed 5-6 times with ice-cold dH_2_O (4K RCF, 4 °C, 10 minutes), and resuspended in fresh buffer (50 mM Tris-SO_4_ pH 7.5, 50 mM NaCl), supplemented with 1 mg.mL^-1^ lysozyme (Roche diagnostics), 0.01 mg.mL^-1^ DNase I (Sigma-Aldrich), 5 mM MgCl_2_, and 1 cOmplete™, Mini protease inhibitor cocktail tablet (Roche diagnostics). This enzymatic treatment was carried out at 4 °C for 45 minutes with gentle agitation primarily to degrade the peptidoglycan cell wall and cortex, before mechanical disruption of the spores by 4-5 passages through a cell disrupter (Constant Systems Ltd) at 30 KPSI to fragment the degraded cortex/membranes and release the core contents. Next, the spore lysate was clarified by centrifugation at 50K RCF, 4 °C for 20 minutes to remove unbroken spores and integument fragments, and the supernatant was ultracentrifuged at 150K RCF, 4 °C overnight (Beckman). The spore membranes were obtained as a red-orange pellet while the supernatant had a yellow colouration. The isolated membranes were washed twice using buffer containing 50 mM Tris-SO_4_ pH 7.5, 50 mM NaCl (150K RCF, 4 °C, 1 hour), homogenised using a Dounce homogeniser (Kimble), aliquoted and stored at −70 °C. The supernatant was syringe-filtered through a 0.22 μm membrane (Sartorius) and concentrated down from ∼50 mL to 5 mL using a 30 kDa MWCO centrifugal concentrator (Sartorius), then aliquoted and stored at −70 °C.

Membranes from *B. megaterium* late-exponential phase cells (16 hours of growth in LB broth medium, 30 °C, 225 rpm shaking incubation) were isolated using a similar method: cell lysis in the presence of 1 mg.mL^-1^ lysozyme and 0.01 mg.mL^-1^ DNase I, 5 mM MgCl_2_ and 1 protease inhibitor tablet by passage through a cell disruptor twice at 30 KPSI. The lysate was centrifuged (10 min, 10K RCF, 4 °C) four times to pellet any debris and the membranes isolated from the supernatant by ultracentrifugation (150K RCF, 4 °C, 1 hr). The isolated membranes were washed twice (1 hr, 150K RCF, 4 °C), aliquoted and stored at −70 °C.

The protein concentration was measured using the Bicinchoninic acid (BCA) protein assay kit (Sigma-Aldrich) following the recommended protocol. BSA (2 mg.mL^-1^) was used to make standards and a dilution series was prepared for the membrane preparations (1/10, 1/20, 1/40, 1/80, 1/160, 1/320, 1/640 and 1/1280). 25 μL standards/sample dilutions were pipetted in triplicate into a 96-well plate followed by 200 μL BCA reagent added to each well. The plate was then incubated at 37 °C for 30 mins. Absorbance was measured at 562 nm in the SpectraMax ABS Plus microplate reader (Molecular Devices) and the data analysed using a predefined protocol in Softmax Pro 7.1.

#### Remission haem spectroscopy and oxygen consumption measurements

Remission haem spectroscopy experiments on cells, spores, and isolated membranes were performed with the prototype of the Iberius Cell Spectroscopy System (CellSpex Ltd) which consisted of a bioenergetic chamber, a single-channel spectroscopy system and an oxygenation system. The bioenergetic chamber had a sample volume of 5 mL. 3.5 mL of *B. megaterium* cells grown in 5 mL LB medium to an O.D. of ∼5.4 (exponential phase, incubated for 12 hours) were diluted with 1.5 mL fresh LB medium to achieve an O.D. of 4. Isolated membranes from cells and spores were diluted in the buffer containing 10 mM Tris-SO_4_ pH 7.3 and 250 mM sucrose such that the final protein concentration was 0.25 mg.mL^-1^. Histodenz-purified and heat-shocked spores were resuspended in 5 mL 50 mM potassium phosphate pH 7.5 buffer to obtain an O.D. of 6. A white LED light source, Luxeon CZ 4000K-90 (Lumileds) used at a current of 200-350 mA, illuminated the samples in a quartz crucible. The back-scattered light was collected in remission geometry, passed through a spectrograph (iHR320, Horiba) equipped with a 300g/mm grating blazed at 500 nm, and complete spectra between 480 and 760 nm were collected on a CCD camera (Andor Technology). The slits were set to 100 μm to give a spectral resolution of 1 nm. The O_2_ optode relies on the phosphorescence half-life of a platinum-porphyrin compound to measure O_2_ concentration (Lee and Okura, 1997). In some experiments, a blend of N_2_/O_2_ was delivered through silicone tubing (Braintree Scientific. Inc.) submerged in the samples to maintain a constant O_2_ concentration throughout (Kim et al., 2011). O_2_ consumption rate was calculated from the difference between the O_2_ delivery to the sample (where tubing was used) and the rate at which the O_2_ concentration in the sample changed during the experiment (Kim et al., 2012; Rocha and Springett, 2019).

To prepare the device for an experiment, a wavelength calibration for the CCD was performed using mercury emission lines (546.074 nm, 576.960 nm, and 579.070 nm), and the optode was calibrated by measuring O_2_ dissolved in the experimental buffer exposed to air at the desired temperature vs. the zero-point achieved by the addition of the strong reducing agent sodium dithionite (Na_2_S_2_O_4_). Then, with the sample buffer in the crucible, tubing in place, and the chamber sealed, the intensity of the LED light source was calibrated (serving as a blank measurement). Following the addition of the cell/isolated membranes/spore samples, they were allowed to equilibrate for 10-15 mins in the re-sealed chamber. The experiments were initiated by the addition of a germinant/NADH/KCN via a small injection port present in the plunger seal at t=0 min and one spectrum was recorded every 20 ms in two phases (each phase is 10 ms long). The device was controlled using the accompanying Palencia software (CellSpex Ltd.).

The phosphorescent light at 650 nm from the O_2_ optode can interfere with attenuance measurements in this wavelength range. The interference was removed using two-phase time multiplexing: the samples were illuminated with a white LED in the first phase but not the second. The spectrum from the second phase was then subtracted from that of the first phase to remove the signal from the O_2_ optode which was present in both phases. Each phase was 10 ms and spectra from 25 pairs of phases were averaged to give a temporal resolution of 500 ms.

The spectra were analysed and manipulated (generation of averaged difference spectra, scaling, Savitzky-Golay smoothing, offsetting) in the Gerona analysis software package (CellSpex Ltd.). The difference spectrum at a given time point T_p_ was generated by averaging the 120 spectra recorded over a 1-min interval preceding the T_p_. From this, the spectrum before glucose/CN^-^ addition was subtracted, and the resulting spectra were fitted to a linear regression to generate the difference spectra shown. Where the Savitzky-Golay smoothing function (Savitzky and Golay, 1964) was used, the half width was 0.5 nm and the order was 1. Decomposition of the haem attenuance spectra was performed using the following equation implemented in Gerona as the ‘FIT:NIR’ model:

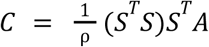

where C is the unknown column matrix containing the concentration of each component, ρ is the differential pathlength, A is the column matrix containing the observed attenuance at each wavelength, and S is the known matrix containing the specific absorbance of each haem centre at each wavelength (Blaza et al., 2014; Kim et al., 2011; Shinkarev et al., 2006). Model spectra for the fitting template were taken from various sources, wavelength-shifted to get the desired peak absorbance, and normalised to 1. The spectra used to generate the cyt *c*-550, *c*-555, *b*-558, *b*-565, and *b*-571 model spectra were originally measured using beef heart cyt *bc*_1_ complex. The Cyt *a*-580 spectrum was obtained from reference (Lauraeus et al., 1993)], and is the spectrum of the fully reduced *B. subtilis aa*_3_-600 with O_2_ at a high pH after 120 mins. Cyt *a*-600 is the haem *a* difference spectrum taken from reference (Liao and Palmer, 1996). The baseline components used were α = 1/x^2^, β = (x-585), ɣ = (x-585)^2^ and δ = 1, where 585 nm is the midpoint of the fitting range (520-650 nm).

#### Spore rehydration and outgrowth assays

Spore germination and outgrowth were assessed by monitoring the change in their initial absorbance at 600 nm. Histodenz-purified *B. megaterium* spores were heat shocked either at 60 ⁰C for 10 mins in a water bath or at 70 ⁰C for 20 mins in a heating block. Histodenz-purified *B. subtilis* spores were heat shocked at 70 ⁰C for 30 minutes in a heating block. The heat-shocked spores were either cooled on ice or washed once (4K RCF and 18K RCF for *B. megaterium* and *B. subtilis* spores respectively, 4 °C, 10 minutes) and resuspended in fresh 50 mM potassium phosphate pH 7.5 buffer. In a 96-well plate with the germinants/nutrient broth/KCN and combinations thereof already present in triplicates, the spore suspension was added to start the assay such that the initial absorbance was 0.6-0.8 in a 200 μL volume. Absorbance measurements at 600 nm were then started immediately in a SpectraMax ABS Plus microplate reader (Molecular Devices) or a Tecan Infinite-200 series monochromatic plate reader fitted with a 600-nm photometric filter. They were set at 30 ⁰C and 37 ⁰C, with a read interval of 10 secs and 30 secs for *B. megaterium* and *B. subtilis* spores respectively. Where specified, assays were performed either with or without orbital shaking for 10 seconds between absorbance readings. Germination/outgrowth assays under anaerobiosis were carried out in an anaerobic/dry glove box system (Belle Technology UK Ltd) maintained at 2.5 ppm (0.00025%) O_2_ with a SpectraMax ABS Plus microplate reader inside. The data were imported from the microplate reader software SoftMax Pro 7.1 and i-control 2.0 into Microsoft Excel and the absorbance values across the technical replicates was averaged. Absorbance measured at t=0 min (t_0_) was taken as the initial absorbance, and the percentage loss of initial absorbance was calculated for all subsequent time points. The % initial A_600_ values were plotted against time to give rehydration and outgrowth curves. The rate of rehydration was given by the first derivative of % loss (t_p_) values. This was calculated using a window of 11 data points and the function SLOPE which returned the slope of the line at the centre of the window as the window moved to the next data point. The first derivative (% loss of initial A_600_/min) was then plotted against time. Experiments were conducted with at least two biological replicates (independently prepared spore batches).

#### Glucose consumption assays

Glucose consumption of germinating *B. megaterium* spores was inferred from the enzymatic quantification of glucose remaining in the germination medium. Pellets of heat-shocked spores were resuspended in 285 μL 50 mM potassium phosphate pH 7.5 buffer and transferred to a 2 mL microcentrifuge tube. All tubes were moved to a heating block set at 30 ⁰C, 300 rpm shaking, with their lids open. Germination was initiated with the addition of 15 μL 20 mM glucose (final concentration of 1 mM in 300 μL volume) and the spores (O.D. of 17) were incubated for the stipulated period (t = 0, 2, 4, 6, 8, 10, 12, 14, 16 mins) after which they were quickly relocated to a heating block set at 100 ⁰C for 10 mins to stop germination, then put in ice until all samples were ready. The tubes were then centrifuged at 16K RCF, 4 ⁰C for 15 mins. The supernatants were transferred to fresh tubes and centrifuged again to remove all the spore debris. The D-Mannose/D-Fructose/D-Glucose assay kit (Megazyme Ltd) was used for the quantification of glucose in these supernatants along with the 1 mM glucose control (without any spores) with minor modifications to the recommended kit protocol. Stoichiometric amounts of NADPH were formed in a 126 μL reaction volume containing 50 μL of each sample. This was performed in triplicates for each time point and the control. The triplicate values from the glucose assay were averaged for each time point and the control, and the amount of glucose consumed (μM) by t minutes was calculated and normalised for O.D. of 17. Glucose consumed (μM/O.D.) values were plotted against time (mins), and a linear regression was fitted in the 6-16 minute region of the curve to get an average rate of glucose consumption. Experiments were conducted with at least two biological replicates (independently prepared spore batches).

#### Hydrogen peroxide production assay

Hydrogen peroxide released by spores germinated with 1 mM/4 mM glucose was measured using an Amplex Red assay kit (Invitrogen, ThermoFisher Scientific). 200 μL reactions were set up in duplicate with heat-activated spores (1 O.D.), 0.5 U.mL^-1^ horseradish peroxidase (HRP) and 50 μM Amplex Red reagent in a 96-well flat-bottom black microplate (Nunc, ThermoFisher Scientific). The negative control reaction, also used for background subtraction during data analysis, contained spores, HRP and Amplex Red but no glucose. The reagent injectors present in the CLARIOstar microplate reader (BMG Labtech) were used to dispense 10 μL of the 20 mM/80 mM glucose stocks to initiate germination, which was immediately following by double orbital shaking at 300 rpm for 30 s, then fluorescence measurements every 15 secs for 30 mins at 30 ⁰C. The excitation/emission wavelengths used were (545-20)/(600-40) which are preset for the reaction product resorufin in the control software, number of flashes/well = 20, gain = 744 and focal height = 8.2 mm. H_2_O_2_ standards were prepared by serial dilution and reactions were set up in triplicate containing the standard, 50 μM Amplex Red and 0.2 U.mL^-1^ HRP. A single-point measurement using the same optic settings was recorded in the microplate reader, and the blank-subtracted fluorescence values were used to plot a standard curve. The blank-subtracted fluorescence values were used to calculate the H_2_O_2_ concentration (nM). The rate of H_2_O_2_ release was given by the first derivative calculated using a window of 5 data points. This first derivative (nM/min) was then plotted against time. Experiments were conducted with at least two biological replicates (independently prepared spore batches).

#### BN-PAGE and LC/MS analysis

BN-PAGE analyses were performed using the well-established NativePAGE Novex Bis-Tris Gel System (Invitrogen, ThermoFisher Scientific). The spore membranes at a protein concentration of 4 mg.mL^-1^ were solubilised with 1% n-dodecyl β-D-maltoside (DDM, GLYCON Biochemicals GmbH) with gentle agitation for 1 hr at 4 ⁰C. The extracted proteins were clarified twice at 16K RCF, 4 ⁰C for 15 mins. The sample was resolved using a pre-cast 3-12% Bis-Tris gradient gel, along with the NativeMark Unstained Protein Standard in a 4 °C cold room at 150 V for the first 60 minutes and at 250 V for the last 30-45 minutes. The gel was later rinsed with dH2O, and fixed for 10 mins in a mixture of 50% ethanol and 10% acetic acid on a gel rocker. After thorough destaining in dH2O, the gel was submerged in the staining solution containing Coomassie brilliant blue G-250 dye and HCl, heated, and left to stain overnight on the gel rocker – this was needed to stain the low abundance membrane proteins for maximum contrast. The following morning, the gel was destained again. Once sufficiently destained, the gel was imaged. Bands were cut out for analysis by the Metabolomics & Proteomics Lab, Technology Facility, Department of Biology, University of York using Liquid Chromatography tandem Mass Spectrometry (LC/MS). A Waters mClass UPLC connected to an Orbitrap Fusion Tribrid mass spectrometer was used for data acquisition, and Progenesis QI was used for chromatographic alignment and peak picking. The protein hits in each BN-PAGE band were then sorted (largest to smallest) based on their peak areas.

#### Phase contrast microscopy

Microscopic examination was performed in tandem with the outgrowth assay. 2 μL of the spore sample from the appropriate well before and during the outgrowth assay were air dried on a glass slide. A cover slip was placed on the nearly dried drop and pressed down to expel air pockets. An Olympus BX53 microscope fitted with a QImaging Retiga 2000R CCD camera microscope, controlled with the software Q-Capture Pro 7 was used to obtain phase contrast images using a 100x objective lens.

### Supplemental methods

#### NADH oxidation assays

The NADH oxidation rates for the membrane preparations were measured spectrophotometrically. The reaction buffer used was 10 mM Tris-SO_4_ pH 7.3 and 250 mM sucrose in a reaction volume of 200 μL in a flat-bottomed 96-well plate containing 200 μM NADH and 0.05 or 0.1 mg.mL^-1^ protein. The volume of the buffer was adjusted accordingly when 1 mM KCN was added. Data were recorded at 340-380 nm every 7 secs for 20 mins in the SpectraMax ABS Plus microplate reader maintained at 30 °C. Rates for the linear region of the oxidation curves were calculated in the software Softmax Pro 7.1. ε_340-380nm_ = 4810 M^-1^cm^-1^ [Birrell et al 2009] was used for the calculations.

#### Chromatographic enrichment of GDH activity

The supernatant obtained after overnight ultracentrifugation of the spore lysate contained soluble proteins present in the core spore. This was syringe-filtered through a 0.22 μm membrane and concentrated down from ∼50 mL to 5 mL using a 30 kDa MWCO centricon and stored at −70 °C. 500 μL of the whole spore soluble fraction was first subjected to SEC (Superdex 200 Increase 10/300 GL, 8 °C), then AEX (HiTrap Q FF, 21 °C) and lastly to SEC again (Superdex 200 Increase 3.2/300 GL, 4 °C). Following each of these chromatographic steps, the resulting fractions were screened for GDH activity using a spectrophotometric assay at 30 °C in a 96-well plate format. 50/5 μL of each SEC fraction was added in a 200 μL reaction also containing 100 μM NAD^+^ and 1 mM glucose. These reactions were performed in a 50 mM Tris-SO_4_ pH 7.5, 50 mM NaCl buffer and absorbance at 340-380 nm was measured every 15 secs or 6-7 secs for 30 mins. The protein concentration of SEC/AEX fractions was also measured with single-point readings using a Bicinchoninic acid (BCA) protein assay kit (Sigma-Aldrich) following the recommended protocol. Dilution factors of 2.5 (after SEC I), 25 (after AEX) and 12.5 (after SEC II) were used – 25 μL of the diluted samples was used for assay as described, and the value obtained was multiplied by the dilution factor to get the protein concentrations of the SEC/AEX fractions. These rates were then plotted against the corresponding fraction number to visualise regions of enriched GDH activity. The fraction with the highest GDH activity was analysed by LC/MS as described in the main text.

